# Engineered dsRNA-protein nanoparticles for effective long-distance transport, delivery and gene silencing in plants

**DOI:** 10.1101/2023.10.12.562099

**Authors:** Huayu Sun, Ankarao Kalluri, Dan Tang, Jingwen Ding, Longmei Zhai, Xianbin Gu, Yanjun Li, Huseyin Yer, Xiaohan Yang, Gerald A. Tuskan, Zhanao Deng, Frederick G. Gmitter, Hui Duan, Challa Kumar, Yi Li

## Abstract

Long-distance transport of exogenous biologically active RNA molecules in higher plants has not been reported. Here, we report that cationized bovine serum albumin (cBSA) avidly binds double-stranded beta-glucuronidase RNA (dsGUS RNA) to form nucleic acid-protein nanocomplexes. Using tobacco and poplar plants, we have shown effective uptake and long-distance transport of cBSA/dsGUS RNA nanocomplexes from basal ends of leaf petioles to leaf blades as well as from basal ends of shoots to their apexes and apical leaves. We have further demonstrated that the cBSA/dsGUS RNA nanocomplexes are highly effective in silencing both conditionally inducible *DR5-GUS* gene and constitutively active *35S-GUS* gene in leaf, shoot and shoot meristem tissues. This cBSA/dsRNA delivery technology may provide a convenient, fast, and inexpensive tool for characterizing gene functions in plants, and potentially for *in planta* gene-editing.

## Introduction

Exogenous RNA application to plants may provide a powerful tool in characterizing gene functions and improving agricultural crop productivity (Das & Sherif, 2020). Exogenously applied naked RNAs were reported effective in gene silencing in plants (Li *et al*., 2015; Dubrovina *et al*., 2019; Molesini *et al*., 2022), but a number of more recent studies have showed that naked RNAs are not effective in gene silencing in plants (Dalakouras *et al*., 2016; Zhang *et al*., 2019; Dalakouras *et al*., 2020; Demirer *et al*., 2020; Zhang *et al*., 2021; Zhang *et al*., 2022; Yong *et al*., 2022). While nanoparticle-RNA complexes have been shown to be highly effective to silence gene expression in living plant cells (Torney et al., 2007; Jiang *et al*., 2014; Ocsoy *et al*., 2018; Zhang *et al*., 2019; Demirer *et al*., 2020; Zhang *et al*., 2021; Hu & Xianyu *et al*., 2021; Azeem *et al*., 2021; Fiol *et al*., 2021; Demirer *et al*., 2021; Zhang *et al*., 2022; Li *et al*., 2022; Yong *et al*., 2022; Molesini *et al*., 2022) but they are relatively inefficient for long-distance transport within the plant. Zhang *et al*. (2019 and 2022) and Demirer *et al*. (2019) reported that nanoparticle complexes diffused only about 60 μm in the *z*-direction and 3 cm in the *x*-*y* direction in the leaves from the infiltrated sites. While there have been reports on the long-distance transport of nanoparticle-RNA in plants, these studies document the silencing effects on genes in fungal pathogens, insect pests, or viruses that infect plants, rather than on endogenous plant genes. (Li *et al*., 2015; Mitter *et al*., 2017; Qiao *et al*., 2023). Therefore, achieving long-distance transport of nanoparticle-RNA complexes is highly desirable, as it would enable systemic gene silencing effects to be obtained, even when double-stranded RNA molecules are applied locally. Here, we report the development of cationized BSA (cBSA) and dsRNA nano-complexes and their long distance-transport and gene silencing activities in higher plants.

## Results

### Chemical modification of cationized BSA and dsRNA nanocomplexes and their transport within the plant

We systematically adjusted the net charge on BSA, for capturing anionic dsRNA as a nanocomplex, by conjugating the COOH groups of BSA with the amine groups of ethylene diamine (EDA) via carbodiimide chemistry (Fig. S1, middle row). The degree of conjugation and net charge on the protein were optimized by adjusting the mole ratios of protein, EDA and the carbodiimide condensing agent (EDC, Fig. S2). For example, the modified protein indicated a high positive charge, moved toward the negative electrode in agarose gel (Fig. S1A, Lane 2) while there was little or no crosslinking (SDS PAGE, Fig. S1B, lane 3, Fig. S3). The reaction conditions were optimized for 100% conversion, without any residual unmodified BSA or protein crosslinking. The cationized BSA (cBSA) with a high positive charge (Fig. S1A, Lane 2) was purified by dialysis and examined by Zeta potential titrations, which indicated a net charge of +23 mV and an isoelectric point of 9.1 (Fig. S1C). The UV circular dichroism (CD) spectra showed that its secondary structure is like that of the native protein (Fig. S1D), while dynamic light scattering (DLS) data (Fig. S1E) indicated a molecular size of 6 nm, further confirming that there has been no protein crosslinking.

The fully characterized cBSA (20 µM, pH 7.2) was then examined to form nanocomplexes with dsGUS RNA (20 µM, pH 7.2) (Fig. S1 middle row) by band shift assay in agarose gel electrophoresis (Fig. S1F, Fig. S4). A mixture of 1:1 mole ratio of cBSA and dsGUS RNA (RNA band stained with SYBR safe dye) showed a significant shift in the band position (Fig. S1F, lane 4), compared to the control lanes (Fig. S1F, Lanes 1-3). The absence of free protein or free dsRNA in this mixture is also evident when stained with Coomassie Blue (Fig. S1F).

The nanocomplex formation was confirmed by *Zeta* potential titrations (Fig. S1G), CD spectra (Fig. S1H) and DLS (Fig. S1I). CD indicated considerable distortion of both the protein secondary structure (peaks at 210 and 222 nm) as well as that of the dsGUS RNA (270 nm band) (Fig. S1H), while DLS indicated a small increase in size from 6 to 8 nm (major fraction) and had a minor fraction of 34 nm. Moreover, cBSA was found to bind dsGUS RNA at 2:1 to 5:1 mole ratios (Fig. S4), but these complexes had much larger particle sizes (40 - 600 nm) and thus were not further examined. Therefore, the smaller 1:1 complex of cBSA/dsGUS RNA was selected to explore gene silencing in plants.

### Systemic silencing effects of cBSA/dsGUS RNA nanocomplexes on auxin-induced expression of the *DR5-GUS* gene

An auxin-inducible synthetic promoter, *DR5* (Ulmasov *et al*., 1997), derived from a soybean *GH3* promoter (Hagen *et al*., 1991; Li *et al*., 1999), was used to drive the expression of the GUS reporter gene in tobacco (*Nicotiana tabacum*). The *DR5-GUS* transgenic tobacco leaf discs were incubated with cBSA/dsGUS RNA at concentrations of 0.01, 0.1, 1, or 10 x 10-6 M for 24 hours. Subsequently, they were treated with auxin (3 x 10-5 M NAA) for 18 hours before histochemical staining of GUS activity. The results of histochemical staining showed that a concentration of 1 x 10-6 M cBSA/dsGUS RNA was sufficient to achieve effective gene silencing. This concentration aligns with the physiological concentrations of plant hormones or growth regulators applied externally, indicating its effectiveness. Fig. 1A illustrates the successful silencing of GUS enzyme activity in representative DR5-GUS transgenic tobacco leaf discs treated with 1 x 10^-6^ M cBSA/dsGUS RNA nanocomplexes. Gene silencing effects were observed in all treated leaf discs. Additionally, we administered cBSA/dsGUS RNA complexes into leaf tissues through petioles (Fig. 1B). Initially, we used intact leaves with petioles for administration and achieved excellent gene silencing effects. However, due to variations in leaf surface areas among individual leaves, we encountered some discrepancies in gene silencing effects. This disparity might be partially attributed to variations in the concentrations of cBSA/dsGUS RNA complexes within leaf tissues due to differences in leaf surface areas. To reduce these variations, we standardized the surface area of tobacco leaves by trimming them (Fig. 1B) to ensure more comparable concentrations of the nanocomplexes across all experimental leaves. Following 72 hours of administration, no evident changes in appearance or health of the experimental leaves were observed. About 0.4 mL of cBSA/dsGUS RNA nanocomplex solution (1 x 10^-6^ M), which was about 6.4 μg of dsGUS RNA in total, was fed into each square centimeter of the experimental leaves during the 72 hours of feeding. We then treated the leaf tissues with NAA (3 x 10^-5^ M) for 18 h before staining for GUS activity. More than 90% of experimental leaves fed with the RNA nanocomplexes displayed reductions in the auxin induced GUS activities while cBSA nanoparticles or naked dsGUS RNA alone did not result in any silencing effects (Fig. 1C). Additionally, qPCR analysis conducted after 24 h of feeding further demonstrated that the mRNA level of the *DR5-GUS* gene in leaves was drastically reduced by cBSA/dsGUS RNA nanocomplexes but not by cBSA nanoparticles or naked dsGUS RNA alone (Fig. 1D). Also, as shown in Fig. 1E and 1F, auxin induced expression of *GH3.1* (Roux & Perrot-Rechenmann, 1997) and *ANN12* (Baucher *et al*., 2011) genes in all leaves was not affected by cBSA/dsGUS RNA nanocomplexes. These results support that the effect of the dsGUS RNA nanocomplexes is gene specific. These results also demonstrate that the cBSA may facilitate long-distance transport of dsGUS RNA from basal ends of petioles to leaf blades, and the silencing effect of the cBSA/dsGUS RNA nanocomplexes is systemic, highly effective and gene-specific.

**Fig. 1.**
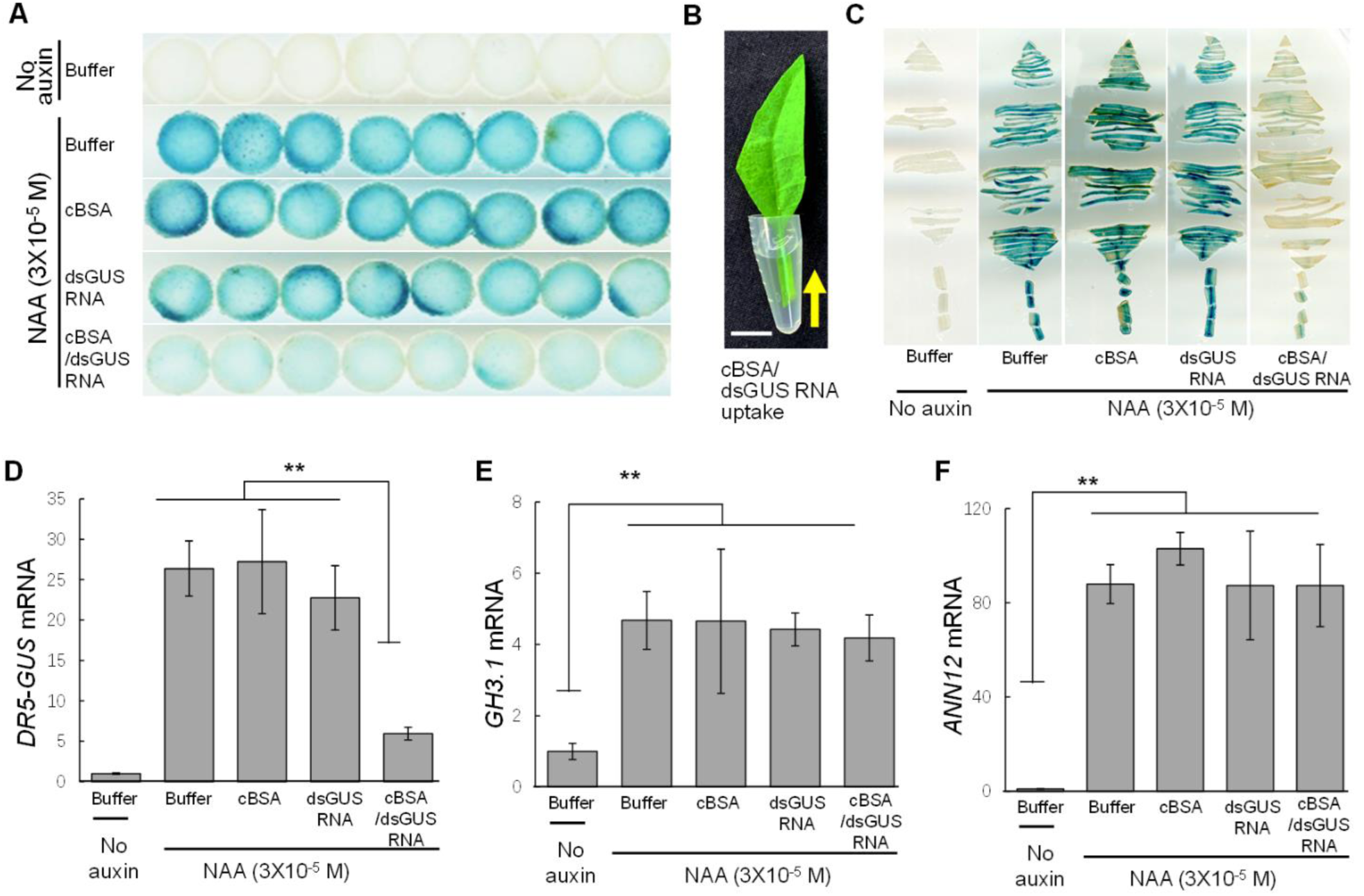
Long-distance transport of cBSA/dsGUS RNA nanocomplexes and their silencing effects on expression of auxin inducible *DR5-GUS* gene in tobacco leaves. (**A**) Histochemical staining of GUS activity shows that cBSA/dsGUS RNA nanocomplexes effectively reduced auxin inducible GUS activity in the *DR5-GUS* transgenic tobacco leaf discs. The leaf discs were incubated in cBSA/dsGUS RNA solution (1 x 10^-6^ M) for 24 h and then treated with auxin (3 x 10^-5^ M NAA) for 18 h prior to histochemical staining of GUS activity. (**B**) cBSA/dsGUS RNA nanocomplexes were fed into leaf tissues from basal ends of leaf petioles. The feeding lasted for 72 h and each cm^2^ of leaf tissues received about 0.4 mL solution of cBSA/dsGUS RNA nanocomplexes (1 x 10^-6^ M). Scale bar, 0.5 cm. (**C**) Histochemical staining of GUS activity of *DR5-GUS* transgenic tobacco leaves shows BSA/dsGUS RNA nanocomplexes effectively reduced GUS activity. (**D**-**F**), qPCR analysis of expression of the *DR5-GUS* gene (**D**), two auxin inducible genes *GH3.1* (**E**) and *NtANN12* (**F**) in the treated leaves shows the GUS transcript was drastically reduced by cBSA/dsGUS RNA nanocomplexes while expression of the *GH3.1* and *NtANN12* was not affected. Elongation Factor 1α gene (*NtEF1α*), a housekeeping gene, was used for normalizing the expression levels of all genes. ** indicates significant differences at p≤0.01 by ANOVA. The data presented are means±AVEDEV, which have been calculated from a minimum of six biological replicates.

We also fed the cBSA/dsGUS RNA nanocomplexes to the basal ends of detached tobacco shoots (Fig. 2A) and observed silencing effects on auxin induced expression of the *DR5-GUS* gene in shoot apical meristem tissues (marked **a** in Fig. 2A) and apical leaves (marked **b** in Fig. 2A). Tobacco shoots, along with their leaves, were fed with a solution containing 1 x 10^-6^ M of cBSA/dsGUS RNA for 72 hours. At the end of feeding, the leaves of the shoots were removed and the bare shoots were longitudinally split into two halves to ensure uniform auxin inducibility during the subsequent auxin treatment. Fig. 2B shows that the auxin induced *DR5-GUS* expression in 10 out of 12 shoot and shoot apex tissues was drastically reduced by the cBSA/dsGUS RNA complexes but not by the cBSA or dsGUS RNA alone. Fig. 2C shows qPCR results that cBSA/dsGUS RNA effectively silenced the auxin induced expression of the *DR5-GUS* gene at the mRNA level, with 85% and 65% reduction in the leaf and shoot meristem tissues, respectively, after 24 h of feeding. Again, as shown in Fig. 2D, the auxin induced expression of the *GH3.1* and *ANN12* genes were not affected by the cBSA/dsGUS RNA nanocomplexes in all shoot tissues including and shoot apex tissues.

**Fig. 2.**
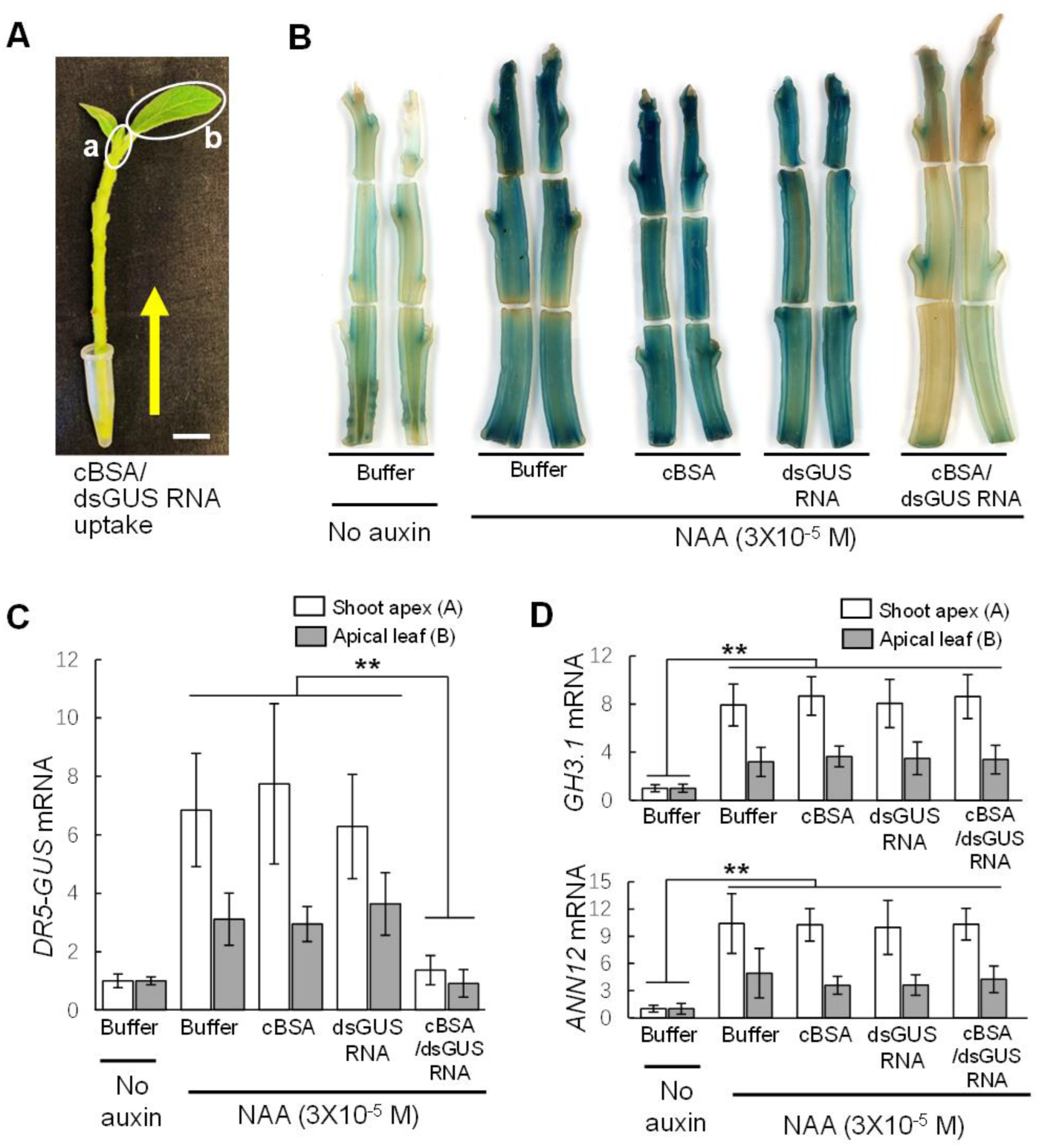
Long-distance transport of cBSA/dsGUS RNA nanocomplexes and their silencing effects on auxin induced expression of the *DR5-GUS* gene in tobacco shoots. (**A**) cBSA/dsGUS RNA nanocomplexes were fed into about 10 cm-long shoots from their basal ends. The feeding lasted for 72 h and a total of about 10 mL solution of cBSA/dsGUS RNA (1 x 10^-6^ M) was fed into each shoot. Scale bar, 1.0 cm. (**B**) Histochemical staining of GUS activities in *DR5-GUS* transgenic tobacco shoots fed with buffer, cBSA nanoparticles, naked dsGUS RNA, and cBSA/dsGUS RNA nanocomplexes. The treated shoots were split longitudinally into two halves after 72 h feeding and treated with auxin (3 x 10^-5^ M NAA) for 18 h before histochemical staining of GUS activity, showing that cBSA/dsGUS RNA complexes effectively silenced the *DR5-GUS* gene expression. (**C**-**D**) qPCR analysis of expression of the *DR5-GUS* (**C**) and auxin inducible *GH3.1* and *ANN12* (**D**) genes in shoot apical tissues (marked **a** in Fig. 2**A**) and leaves (marked **b** in Fig. 2**A**), demonstrating that cBSA/dsGUS RNA nano-complexes silenced *DR5-GUS* gene at the mRNA level and also the effects are specific. Expression levels of *NtEF1α* was used for normalizing the expression levels of all genes assayed. The white and gray filled bar graphs represent apical meristem tissues (marked **a** in Fig. 2**A**) and leaf tissues (marked **b** in Fig. 2**A**), respectively. ** indicates significant differences at p≤0.01 by ANOVA. The data presented are means±AVEDEV, which have been calculated from a minimum of three biological replicates.

### Systemic silencing effects of the cBSA/dsGUS RNA nanocomplexes on constitutively active genes

To explore efficiencies of the cBSA/RNA nanocomplexes on silencing of constitutively active genes, leaves of *35S-GUS* transgenic tobacco and poplar plants were detached and fed with the cBSA/dsGUS RNA complexes (1 x 10^-6^ M) through petioles (Fig. 3A and Fig. 3B). As shown in Fig. 3A, we observed 60% and 70% of the constitutively active *35S-GUS* gene in all experimental leaves (6 out of 6) of poplar and tobacco, respectively. The cBSA nanoparticles or naked dsGUS RNA had no silencing effects on expression of the *35S-GUS* gene in either plant species. We also administered the cBSA/dsGUS RNA nanocomplexes to the basal ends of *35S-GUS* transgenic poplar shoots (Fig. 3C). Consequently, using a qPCR technique, we detected approximately 60% reduction in the *35S-GUS* mRNA levels in shoot apical/meristem tissues in all experimental poplar shoots. Throughout the feeding period of 72 hours, about 6 ml of cBSA/dsGUS RNA nanocomplexes (about 96 ug of dsGUS RNA in total) was absorbed by each poplar shoot, including its leaves. These results demonstrate that the cBSA/dsGUS RNA nanocomplexes can also silence constitutively active genes after long-distance transport in shoot apex tissues.

**Fig. 3.**
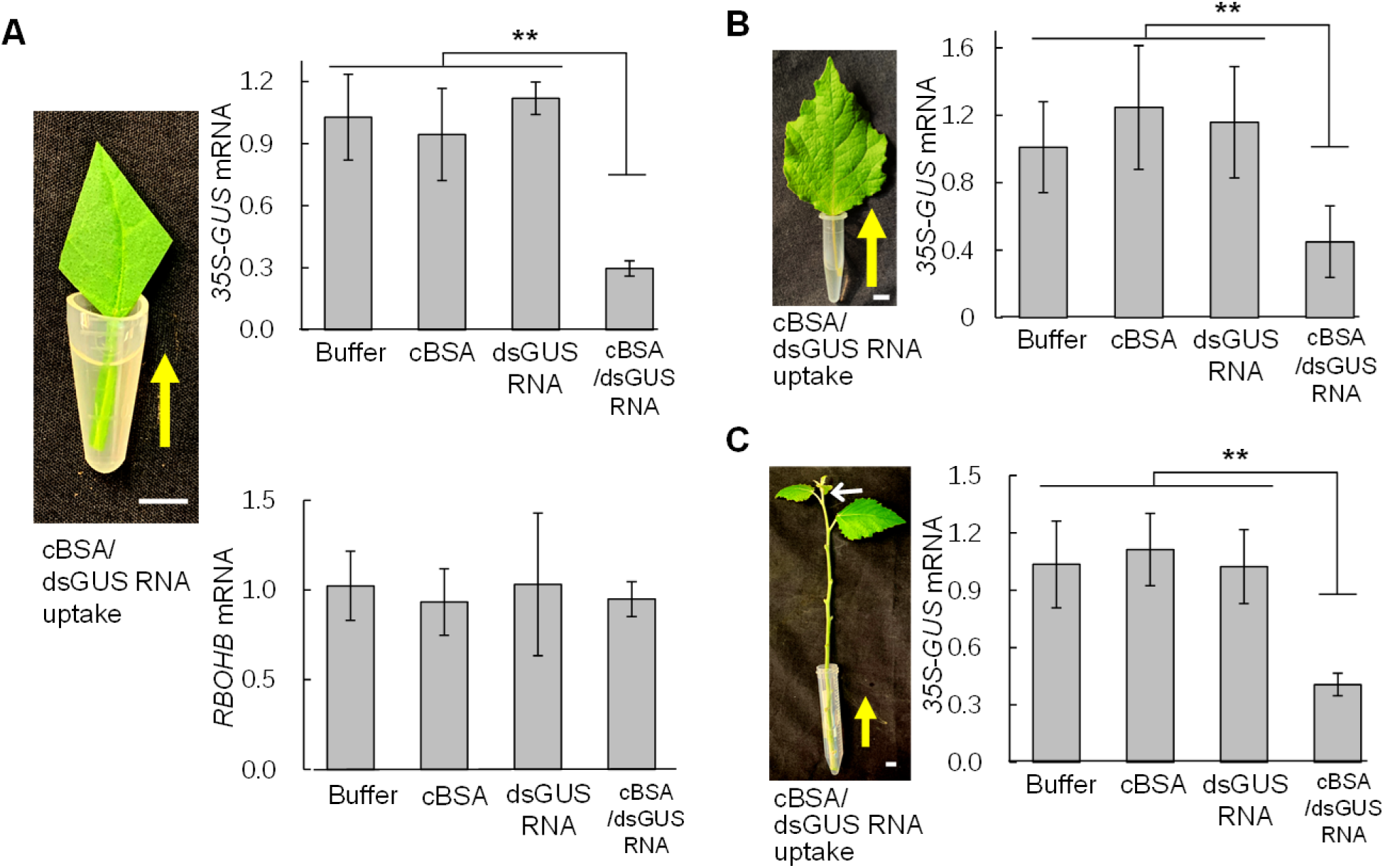
Long-distance transport of cBSA/dsGUS RNA nanocomplexes and their silencing effects on expression of constitutively active *35S-GUS* gene in tobacco and poplar. (**A**) Constitutively active *35S-GUS* gene in tobacco leaves was specifically silenced at the transcript level by cBSA/dsGUS RNA nano-complexes while expression of an oxidative stress inducible *RBOHB* gene was not affected. cBSA/dsGUS RNA nano-complexes were fed from basal ends of leaf petioles for five days and approximately 0.6 mL cBSA/dsGUS RNA nano-complex solution (1 x 10^-6^M) was fed into each square centimeter of the leaves. Scale bars, 1.0 cm. (**B**) Constitutively active *35S-GUS* gene in poplar leaves was specifically silenced at the transcript level by cBSA/dsGUS RNA nano-complexes fed from the basal ends of petioles. The feeding lasted for five days and approximately 0.5 mL nano-dsGUS RNA nano-complexe solution (1 x 10^-6^M) was fed into each square centimeter of the leaves. Scale bars, 1.0 cm. (**C**) Constitutively expressed *35S-GUS* gene in poplar shoot meristem tissues (white arrow pointed) was specifically silenced at the transcript level by cBSA/dsGUS RNA nanoparticle complexes fed from the basal ends of 15 cm-long shoots. The feeding lasted for 6 days and a total of about 6 mL cBSA/dsGUS RNA nano-complexe solution (1 x 10^-6^M) was fed into each shoot. Scale bars, 1.0 cm; For all qPCR analyses, expression levels of *NtEF1α* or *PtEF1α* were used as internal controls for normalizing expression levels of the *35S-GUS* gene. ** indicates significant differences at p≤0.01 by ANOVA. The data presented are means±AVEDEV, which have been calculated from a minimum of three biological replicates.

Further, we have also shown that cBSA/dsPAP2 RNA nanocomplexes is effective to silence expression of the Production of Anthocyanin Pigment 2 (*PAP2*) gene. Feeding the cBSA/dsPAP2 RNA nanocomplexes through petioles of *35S-PAP2* transgenic tobacco, we observed 35% reduction in the *35S-PAP2* expression in all of the feeding leaves (6 out of 6) (Fig. S5). We used oxidative stress inducible *RBOHB* gene (Yoshioka *et al*., 2003; Demirer *et al*., 2019; Zhang *et al*., 2021) to monitor whether cBSA/dsGUS RNA nanocomplexes may cause unintentional stresses to the plant tissues and therefore alter expression of some genes nonspecifically. Fig. 3A shows that cBSA/dsGUS RNA nanocomplexes do not alter *RBOHB* gene expression, indicating that the silencing effects on the *35S-GUS* gene should be gene-specific.

## Discussion

RNA interference (RNAi) technologies have significant potential for enhancing the productivity of agricultural crops. However, the production of RNAi transgenic plants often involves expensive and time-consuming processes. Early studies demonstrated gene silencing in plants using naked RNA, but more recent studies have shown that the application of naked RNA molecules does not result in gene silencing in plants (Dalakouras *et al*., 2016; Zhang *et al*., 2019; Dalakouras *et al*., 2020; Demirer *et al*., 2020; Zhang *et al*., 2021; Zhang *et al*., 2022). We believe that naked dsRNA molecules unlikely enter plant cells due to their negatively charged nature, which hinders their ability to cross the plasma membrane.

Nanoparticles have been successfully utilized to facilitate RNA delivery into plant cells (Zhang *et al*., 2019; Demirer *et al*., 2020; Zhang *et al*., 2021; Jiang *et al*., 2014; Zhang *et al*., 2022). These nano-RNA complexes aid in the delivery of RNA molecules across the plant cell plasma membrane, resulting in the silencing of target genes. However, the silencing effects of these complexes are somewhat limited to the local areas of application (Jiang *et al*., 2014; Zhang *et al*., 2019; Demirer *et al*., 2020; Zhang *et al*., 2021; Zhang *et al*., 2022). Also, some nanoparticles such as LDH nanosheets and artificial nanovesicles have been shown to facilitate the long-distance transport of dsRNA molecules (Li *et al*., 2015; Mitter *et al*., 2017; Qiao *et al*., 2023). However, it is unknown whether these dsRNA-nanocomplexes can penetrate plant cell membrane, as these studies have primarily focused on investigating their impact on gene expression in insects, fungi, or viruses that infected the experimental plants (Li *et al*., 2015; Mitter *et al*., 2017; Qiao *et al*., 2023). As shown here, our cBSA/dsRNA nanocomplexes may present a highly useful and more convenient tool that not only enables the delivery of RNA molecules into plant cells but also facilitates their long-distance transport within the plant and systemic silencing effects of target genes.

Developing nano-RNA delivery technologies that can be applied to plants without the need for external forces is highly desirable. In previous studies, the delivery of nano-RNA or -DNA complexes into leaf tissues often required the use of external forces, such as needle-less syringe-mediated infiltration (Jiang *et al*., 2014; Zhang *et al*., 2019; Demirer *et al*., 2019; Demirer *et al*., 2020; Zhang *et al*., 2021; Zhang *et al*., 2022). Nevertheless, our cBSA/dsRNA nanocomplexes can be easily administered to plants by feeding them through the basal ends of leaf petioles or stems/shoots. Subsequently, they can be transported to various regions of the leaves or shoots without relying on external forces such as high-pressure-mediated infiltration.

The small size of the cBSA/dsRNA nanocomplexes, their strong association and neutral charge provide possible reasons for their efficient long-distance transport and systemic effects in plant tissue. BSA is a globular monomeric protein of molecular mass of 65.4 kDa with 7 nm diameter (Axelsson, 1978). To minimize the nanocomplex size for effective transport in the plant and across cell membranes, we adjusted the charge on cBSA to obtain a soluble RNA complex at 1:1 mole ratio. The cBSA is nontoxic to the plants, highly water soluble, and enabled RNA transport over long distances in the plant. In total, the cationized BSA-based gene delivery platform is facile, bioactive, environmental-friendly, highly water soluble, scalable and inexpensive to be multiplexed for silencing multiple gene targets.

The efficient long-distance transport characteristics of the cBSA/dsRNA nanocomplexes may make the technology versatile for other applications. One potential application is that our cBSA/dsRNA delivery technology may facilitate delivery of gRNA molecules to shoot meristem cells of *Cas9* expressing plants, so that *“in planta”* gene editing may be achieved. For this, gRNA nanocomplexes can be applied to the basal ends of shoots/stems or directly injected into the stem tissues of a Cas9 expressing platform plant. The cBSA nanoparticles can expedite the transport of gRNAs to the shoot meristem cells, where gene editing may occur. Consequently, emerging shoots originating from these meristem cells can be gene-edited. This method holds promise due to its simplicity and efficiency, eliminating the need for tissue culture, plant regeneration, or repeated genetic transformations for every gRNA introduced.

Another potential application is that the cBSA/dsRNA nanocomplexes might be used to transiently silence target genes in large-scale, field-grown plants, particularly orchard trees, through trunk injection (Archer *et al*., 2022) for enhancing productivity. For example, silencing of the farnesyltransferase, *RACK1*, *OsGRXS17* and *BrDST71* genes can enhance plant’s drought tolerance (Wang *et al*., 2009; Li *et al*., 2009; Hu *et al*., 2017; Park *et al*., 2018). Transient silencing of these genes using the cBSA/dsRNA technology may offer a safe, low-cost, non-transgenic approach for enhancing productivities of field-grown crops. Currently, the cost for dsRNA production is less than $0.5/g ($500/kg) with no length limitation, and the cost of the short dsRNA production will be further reduced in future (Taning *et al*., 2020), which can make large-scale field applications of nano-dsRNA complexes economically feasible.

## Conclusions

In summary, we have developed and used environmentally friendly and low-cost cBSA/dsRNA nanocomplexes for efficient delivery and long-distance transport of dsRNA molecules in plants. We have demonstrated that the long-distance transported cBSA/dsRNA nanocomplexes are highly effective to silence both conditionally inducible and constitutively active genes in tobacco and poplar. The cBSA/dsRNA delivery technology described here may provide a convenient, fast and non-transgenic tool to manipulate gene expression in higher plants that may be used for functionally characterizing novel genes, for *in-planta* gene editing, and for enhancing crop productivity under field conditions.

## Materials and methods

### Material Preparation

Bovine serum albumin (BSA) was purchased from Equitech-Bio Inc. (Kerrville, TX). Ethylenediamine (EDA) and N-(3-Dimethylaminopropyl)-N′-ethyl carbodiimide hydrochloride, 98% EDC [1-(3-Dimethylaminopropyl)-3-ethyl carbodiimide hydrochloride] were purchased from Sigma Aldrich. SYBR Safe nucleic acid gel staining dye was obtained from Invitrogen (Fisher Scientific, MA). All chemicals were used directly without any further purification. Dialysis membrane (25 kDa MWCO) was purchased from Spectrum Laboratories, Inc. (Rancho Dominguez, CA). Biology grade agarose was purchased from Hoefer Inc. (Holliston, MA). Two pieces of dsGUS RNA (GUS1 and GUS4), which were fully complementary to the GUS mRNA, were synthesized using the cell-free bioprocessing platform of Genolution (Seoul, South Korea). The GUS1 and GUS4 dsRNA sequences, which have lengths of 126 bp and 127 bp respectively, were combined and then subjected to sonication to produce shorter fragments ranging from 20 bp to 30 bp in length. These fragmented sequences were then used for the further experiments (see Supplementary Table 1 for the sequences of dsGUS RNA).

### Synthesis of the Cationized BSA (cBSA)

cBSA protein nanoparticles were prepared based on a modified version of previously reported procedure (Kumar *et al*., 2015), which is described here. A stock solution of EDA (0.5 M, 50 mL, pH 5.1) was prepared with deionized (DI) water. Next, BSA solution (0.1 g, in 8 mL of 0.1 M phosphate buffer, pH 5.1) was added to 1.5 ml of the EDA stock solution. The mixture was stirred for 15 minutes at room temperature, and then a fresh solution of 0.5 ml EDC (0.1 M, 0.5 mL in DI) was added to the mixture, which was stirred continuously. The final molar ratios of BSA: EDA: EDC were 1: 200: 200 (total of 10 ml of reaction volume), respectively. The mixture was stirred for 4 h at room temperature and the condensation reaction was stopped by adding 1 mL of acetate buffer (4M, pH 4.75) to each reaction mixture. The excess EDA, EDC, and byproducts were removed by dialysis against a 10 mM sodium phosphate buffer at pH 7.2 using a 25,000 Da cutoff dialysis membrane for 2 days. After dialysis, the product yield cBSA was 80% (∼8 mg/mL) as estimated by BSA absorbance at 280 nm and the known extinction coefficient. The net charge on cBSA was confirmed by *zeta* potential measurements described below. The degree of chemical modification was carefully controlled by adjusting the EDA and EDC concentrations such that modified BSA charge varied gradually from a net negative to a net positive value.

### Synthesis of the cBSA/dsGUS RNA nanoparticle soluble complexes

cBSA/dsGUS RNA complex nanoparticles were prepared by adding dsGUS RNA (20 μM, 5 ml) to a cBSA solution (20 μM, 5 ml) at various N/P (nucleic acid and protein) molar ratios followed by immediate gentle mixing. When conducting the bulk and plant feeding experiments, we prepared a dilute cBSA (75 μM, 5 ml, pH 7) solution mixed with dsGUS RNA (75 μM, 5 ml) to make cBSA/dsGUS RNA-NPs with the required molar ratios and the resulting solutions were incubated for 1 day. During this step, the resulting complex solutions were precipitated, then diluted to < 1 µM concentration to solubilize the protein-nucleic acid complexes. The unbound protein or dsGUS RNA were removed by precipitation of the complexes (high concentration >10 μM) followed by centrifugation and redispersal of the nanocomplexes in buffer solution. When performing bulk preparation, cBSA/dsGUS RNA complexes were mixed in a 1:1 molar ratio (1 µM, 10 ml) and the nanocomplexes were equilibrated for 1 day, these complexed nanoparticles were further used in the plant feeding experiments, unless otherwise noted.

### Agarose gel electrophoresis

Agarose gel electrophoresis was performed using a method reported earlier, using a horizontal gel electrophoresis apparatus (Gibco model 200, Life Technologies Inc, MD) and agarose (0.5% w/w) in tris acetate (40 mM, pH 8.2) buffer. Modified cBSA conjugates were loaded with 50% loading buffer (50% v/v glycerol and 0.01% w/w bromophenol blue). Detection of the dsGUS RNA and cBSA/dsGUS RNA complexes complex samples was accomplished through additional staining with SYBR Safe DNA gel staining dye. Samples were dispensed into wells placed in the middle of the gel to allow the protein to migrate towards the negative or positive electrode, based on net surface charge. The formation of cBSA/dsGUS RNA complexes was confirmed by a gel shift assay in a native agarose gel electrophoresis. A potential of 100 V was applied for an appropriate duration, gels were stained overnight with 10% v/v acetic acid, 0.02% w/w Coomassie blue, followed by destaining in 10% v/v acetic acid, overnight.

### Sodium dodecyl sulphate polyacrylamide (SDS-PAGE) gel electrophoresis

Prepared polyacrylamide separating gels (12.5% w/w) and stacking gels (5% w/w) were directly used for SDS-PAGE gel to run the samples. The separating gel was poured into a 1.25 mm mold and allowed to polymerize for 30 minutes (6.5 cm long), after which the stacking gel was poured and similarly allowed to polymerize (1.5 cm long). Samples prepared by combining 10 μL of ∼100 μM BSA-EDA conjugates with 30 μL SDS-PAGE loading buffer (2% w/v SDS, 10% w/w 2-mercaptoethanol), after which the solutions were heated at 90 °C for 2 minutes and 20 μL of each sample solution was loaded per well. Gels were run in a vertical Bio-Rad Mini electrophoresis apparatus at 200 V until the dye front reached the bottom of the gel (∼45 minutes). 800 ml of SDS-PAGE Running gel buffer (25 mM Tris, 192 mM glycine, 3.47 mM (0.1%) SDS, pH 8.3) for single gel run experiment. The gel was stained with Stain A solution (10% v/v acetic acid, 10% v/v isopropanol, and 0.02% Brilliant blue R250) for four hours, followed by stain B solution (20% v/v acetic acid and 0.03 % Brilliant blue R250) for 4 hours. Gels were destained with 10% acetic acid overnight prior to imaging.

### Dynamic light scattering (DLS)

Hydrodynamic radii of dsGUS RNA, cBSA and cBSA/dsGUS RNA complexes were measured using a CoolBatch dynamic light scattering apparatus as reported earlier, using a Precision Detectors (Varian Inc., NJ) with a 0.5 x 0.5 cm^2^ cuvette and a 658 nm excitation laser source at 90° geometry. The samples of the dsGUS RNA, cBSA and the nano-complex (20 μM) were diluted with 1 mL of phosphate buffer (100 mM, pH 7.2) and equilibrated for 300 s at 25 °C at 5 repetitions with 100 accumulations. Precision Elucidate (v1.1.0.9) was used to run the experiment, and Deconvolve (v5.5) was used to process the data.

### *Zeta* potential studies

The surface charge of cBSA and dsGUS RNA as well as complexed nanoparticles was characterized by using zeta potential analyzer. The samples were prepared by mixing 15 µM with 1 mM KCl electrolyte solution. The zeta potential was got from Zeta potential analyzer (Brookhaven zeta plus, Holtsville, NY). The sample volume of 1.6 mL was taken in polystyrene cuvette then electrode for zeta-potential analyzer immersed in the solution and connected to the instrument. The sample’s *zeta* potential calculated by using laser Doppler velocimetry and Smoluchowski fit of the instrument. Each measurement is the average of three runs with triplicates of different samples and the standard deviation calculated.

### Circular dichroism (CD) study

CD spectroscopy was used to monitor the retention of secondary structure, or any conformational changes in dsGUS RNA, cationic BSA (cBSA) and cBSA/dsGUS RNA nanocomplexes. The change in the ellipticity was recorded by using a Jasco 710 spectropolarimeter. Each sample concentration was around 5 µM dissolved in 10 mM phosphate buffer (pH 7.2) to record the ellipticity values. Spectra were obtained using a 1 cm path length quartz cuvette from 300 nm to 200 nm. While collecting the data, the sensitivity of samples was set as 100 mdeg and the data pitch as 1 nm, with continuous scanning mode, 50 nm min^-1^ scanning speed, 1 s response, 1.0 nm bandwidth. The baseline was measured with 10 mM phosphate buffer (pH 7.2) and subtracted from each spectrum. Three scans were used to get the average data then normalized to millimolar concentration of the RNA or protein per unit path length.

### Plant growth

Transgenic *DR5-GUS* (*Nicotiana tabacum*), *35S-GUS* (*N. tabacum* and *Populus alba*×*P. berolinensis* hybrid clone) and *35S-PAP2* (*N. tabacum*) seeds were germinated and seedlings were grown in a substrate consisting of a mixture of soil and vermiculite (3:1). The air relative humidity was approximately 80%. The tobacco seedlings were grown for 3-4 weeks and the poplar seedlings were grown for 5-6 months under a light intensity of 400 mol·m^-2^·s^-1^ before being used for experimentation. The white tobacco seedlings (*Nicotiana tabacum*) with *PDS* gene mutation was cultured in the MS solid medium.

### Nanocomplexes and auxin treatment of plant tissues

The *DR5-GUS* and *35S-GUS* transgenic tobacco leaves were detached to a diamond-shapes with leaf petiole in the middle. Leaves of similar size were used for the feeding experiment, the cBSA/dsGUS RNA complexes were fed through the basal ends of leaf petioles. For poplar leaves, intact young leaves were used, and the nanocomplex feeding experiment was conducted through the basal ends of leaf petioles. To maintain plant growth, 1 x Hoagland medium (Sigma-Aldrich) was added to the feeding solution every two days. Before auxin induction, the leaf tissues were cut into slices. Additionally, leaf disks of 0.8 cm in diameter were cut out from *DR5-GUS* transgenic tobacco leaves using a hole puncher. Subsequently, these leaf disks were immersed in the cBSA/dsGUS RNA (1 x 10^-6^ M) nanocomplex solution with gentle agitation for 24 h before auxin induction with a light intensity of 100 mol·m^-2^·s^-1^.

The *DR5-GUS* tobacco shoots, which were approximately 10 cm long, were detached with two apical young leaves left. The detached shoots were then fed with cBSA/dsGUS RNA complexes from their basal ends. The *35S-GUS* poplar leaves, with the petiole left intact, were directly used in the feeding experiment by inserting the basal ends of leaf petioles into the feeding solution with a light intensity of 100 mol·m^-2^·s^-1^. The *35S-GUS* poplar shoots (∼15-cm long) detached with two apical young leaves left were fed with nanocomplexes from their basal ends. To maintain plant growth, 1 x Hoagland medium (Sigma-Aldrich) was added to the feeding solution every two days. The shoot tissues were split longitudinally into two halves before auxin induction.

For the auxin treatment, all leaf and shoot tissues were incubated in an NAA solution (3 x 10^-5^ M) with gentle agitation for 18 h with a light intensity of 100 mol·m^-2^·s^-1^ before histochemical staining of GUS activity and qPCR.

### Histochemical staining of GUS activity

Histochemical assays for GUS activity was conducted using the method described by Li *et al*. (1999). The tissues were immersed and incubated in the GUS dye solution at 37°C for 12-20 h. Pigments in the tissues were removed through incubation in an increasing ethanol gradient which progressed as follows: 20%, 40%, 60%, 80%, and 95%. After the pigments were fully removed, the tissues were rehydrated serially by 80%, 60%, 40%, 20%, and 0% ethanol before being analyzed photographically.

### qPCR analysis

The qPCR was performed to quantify the expression of target genes in plants using the following commercially available kits: NucleoSpin RNA (MN, Germany) for total RNA extraction, iScript cDNA synthesis kit (Bio-Rad, USA) for reverse transcription of total RNA into cDNA, and iTaq universal SYBR green supermix (BioRad, USA) for qPCR. The target genes for qPCR were *GUS*, *NtANN12*, and *RBOHB*. The expression levels of *NtEF1α* in tobacco (Schmidt & Delaney, 2010) and *PtEF1β* in poplar (Wang *et al*., 2013) were used to normalize the expression levels of each target gene. The sequences of each pair of primers are listed in Supplementary Table 2. All steps followed the guidelines of MIQE (Minimum Information for publication of Quantitative real-time PCR Experiments).

## Data Availability Statement

The data that support the findings of this study are available from the corresponding authors upon request; and the detailed information of primer and dsRNA sequences used in this study is provided in supplementary documents.

## Acknowledgements

This work was supported by the USDA National Institute of Food and Agriculture SCRI (grant no. 2015-70016-23027) and the Connecticut-Storrs Agriculture Experimental Station.

## Author contributions

Yi Li and C.K. conceived the study. Yi Li, C.K., H.S., A.K., X.Y., G.A.T., Z.D. and F.G.G. were involved in the experimental designing. H.S., A.K., D.T., J.D., L.Z., X.G., Yanjun Li, H.Y., L.Z. and X.G. performed the experiments. Yi Li, C.K., H.S. and A.K. conducted the data analysis. Yi Li, C.K., H.S., A.K., X.Y., G.A.T., Z.D. and F.G.G. wrote and edited the manuscript.

## Supporting information

**Fig. S1** Synthesis and characterization of cationized BSA (cBSA) and cBSA/dsGUS RNA nanocomplexes. Schematic illustration of the synthesis of cBSA followed by cBSA/dsGUS RNA nanoparticle complexes (middle row). Cationization of BSA by amidation of its carboxylic groups with ethylenediamine (EDA) using carbodiimide (EDC) chemistry (left hand). The cBSA binds to dsGUS RNA together via intermolecular H-bonding and strong electrostatic interactions (right hand). (**A**) Agarose gel (0.5%) of pure BSA (lane 1) and cBSA (lane 2) protein nanoparticles stained with Coomassie blue. (**B**) The SDS-PAGE of the molecular weight marker (lane 1), unmodified BSA (lane 2), cBSA (lane 3). (**C**) Zeta potential of cBSA as a function of pH (2∼14). The arrow indicates the isoelectric point (PI) at pH 9.1. (**D**) Circular dichroism (CD) spectra of BSA and cBSA. (**E**) the average hydrodynamic diameter of the cBSA was around 7 nm at pH 7. (**F**) cBSA/dsGUS RNA nanocomplex titrations with agarose gels stained with SYBR green dye (left) and stained with Coomassie blue to visualize the protein bands (right). Lane 1: dsGUS RNA, Lane 2: BSA/dsGUS RNA, Lane 3: EDA/dsGUS RNA, Lane 4: cBSA/dsGUS RNA. The Lane 2 and 3 indicates, neither native BSA nor the EDA does not interact with dsGUS RNA, however, Lane 4, cBSA and dsGUS RNA (1:1 mole ratio) shows stronger binding affinity forms ionic nanocomplex. The gels ran at pH 8.2 in 40 mM Tris-acetate buffer. (**G**) Zeta potential of cBSA/dsGUS RNA nanocomplexes with respective their moles ratios. (**H**) Far-UV circular dichroism (CD) spectra of phosphate buffer (10 mM, pH 7.0, gray), dsGUS RNA (5 µM, blue), cBSA (5 µM, black) and cBSA /dsGUS RNA nanocomplex (5 µM; 1:1 mole ratio, orange). (**I**) Average diameter (nm) of the cBSA/dsGUS RNA nanocomplex was around 7 nm at the pH 7 measured through DLS.

**Fig. S2** Agarose gel electrophoresis of BSA and BSA-EDA conjugates stained with Coomassie blue. The samples were spotted at the center of the gel and migrated to up or down depending on their charge. Lane 1: BSA, Lane 2: EDA, Lane 3-7: reaction mixture of BSA, EDA and EDC in the mole ratios as indicated in each lane. Electrophoresis was done at pH 7.0 in 40 mM Tris-acetate buffer.

**Fig. S3** The SDS-PAGE of the BSA-EDA conjugates. Lanes 1 & 6: molecular weight markers; Lane 2: unmodified BSA; Lane 3-6: BSA-EDA conjugates at increasing stochiometries.

**Fig. S4** cBSA titrations with dsGUS RNA as monitored in agarose gel electrophoresis. (**A**) Stained with SYBR green dye to see RNA bands, and (**B**) Stained with Coomassie blue to see protein bands. The numbers above the wells in both gels represent the mole ratios of cBSA to dsGUS RNA, lane 1: dsGUS RNA; lanes 2-7: increasing mole ratio of cBSA to dsGUS RNA; lane 8: cBSA, no RNA. The gel was run at pH 7.0 in 40 mM Tris-acetate buffer.

**Fig. S5** Long-distance transport of cBSA/dsPAP2 RNA nanocomplexes and their silencing effects on expression of constitutively active *35S-PAP2* gene in tobacco. (**A**) cBSA/dsPAP2 RNA nanocomplexes were fed into leaf tissues from basal ends of leaf petioles. The feeding lasted for 72 h and each cm^2^ of leaf tissues received about 0.4 mL solution of cBSA/dsGUS RNA nanocomplexes (1 x 10^-6^ M). Scale bar, 0.5 cm. (**B**-**C**) qPCR analysis of expression of the *35S-PAP2* gene in the treated leaves shows the *PAP2* transcript (**B**) was reduced by cBSA/dsPAP2 RNA nanocomplexes, while expression of an oxidative stress inducible *RBOHB* gene (**C**) was not affected. Elongation Factor 1α gene (*NtEF1α*), a housekeeping gene, was used for normalizing the expression levels of all genes. ** indicates significant differences at p<0.01 by ANOVA. The data presented are means±AVEDEV, which have been calculated from a minimum of six biological replicates.

**Table S1** The sequence information of double-strand GUS RNA. Two pieces of dsGUS RNA (dsGUS1 and dsGUS4), fully complementary to the *DR5-GUS* and *35S-GUS* mRNA in the transgenic tobacco and poplar, were synthesized by Genolution (Seoul, Korea).

**Table S2** The sequence information of primers for qPCR analysis.

